# Population size estimation when multiple samples carrying the risk of misidentification are taken within the same capture occasion from the same individual

**DOI:** 10.1101/2024.06.12.598605

**Authors:** Rémi Fraysse, Rémi Choquet, Roger Pradel

## Abstract

Although non-invasive sampling is increasingly used in capture-recapture (CR) monitoring, it carries a risk of misidentification that, if ignored, causes an overestimation of population size. Models that deal with misidentification have been proposed. However, these models assume that only one sample can be collected per individual at one occasion. This is not true for several monitoring programs based on DNA, for example for those that extract the DNA from faecal samples. The models do not take repeated observations into account, leading to biased estimates.

In this paper, we develop an approach that extends the latent multinomial model (LMM) of Link et al., 2010 using a Poisson distribution to model the number of samplings of the same individual on a given occasion. We then conduct simulations to test how our new model performs. As an illustration, we applied the new Poisson model to a collection of Eurasian otter faeces (Lampa et al., 2015).

Our model yields unbiased estimates of population size when the expected number of samples per individual (*λ*) is sufficiently high: simulations with *λ* ≥ 0.36 and five capture occasions or with *λ* ≥ 0.23 and seven or more occasions. In contrast, when *λ* = 0.11 (corresponding to about 42%, 53% and 62% of the individuals being detected with respectively 5, 7 and 9 occasions), the population size is consistently underestimated. Applying the model to the otter dataset confirms the presence of misidentifications, consistent with the authors’ expectations.

Our findings indicate that repeated observations can be modelled without bias. The application on otters shows that our model is necessary to accurately estimate population size in presence of misidentification and repeated observations.

## Introduction

One question that can be crucial to answer in conservation management is “How many individuals are there in a population?”. To answer that question, capture-recapture models are powerful methods that require previously captured individuals to be recognised. Traditional monitoring protocols use marks posed on individuals upon initial capture. But non-invasive protocols such as DNA-sampling of faeces, hair or feathers can also be used when capture proves difficult. Studies using DNA-sampling are common and have been carried out on bears (Dreher et al., 2007), bobcats (Morin et al., 2018; Ruell et al., 2009), pronghorns (Woodruff et al., 2016), elephants (Laguardia et al., 2021) or otter (Lampa et al., 2015), for example. While non-invasive sampling allows the study of free-ranging, elusive species without the need to capture, handle or, in some cases, even observe them, compared to traditional tagging methods, individual identification is however not straightforward. This method carries a much higher risk of incorrect individual identification (Taberlet et al., 1999). If misidentifications are ignored, then standard models would overestimate population size, by a factor that could reach up to five times depending on conditions (e.g. Creel et al., 2003).

Proposals to reduce misidentifications from DNA samples range from adapted field methods and improved laboratory techniques (Paetkau, 2003; Waits and Paetkau, 2005) to pre-analysis software that helps filter out data likely to contain errors (McKelvey and Schwartz, 2005). In addition, various approaches have been put forward to model misidentifications when estimating population size (Link et al., 2010; Lukacs and Burnham, 2005; Wright et al., 2009; Yoshizaki et al., 2011). Today, the most common practice is still to filter out the DNA samples that have a too small number of reliably amplified loci.

However, discarding data may result in too little data being retained for reliable estimation of the parameters of interest. In such cases, it may be beneficial to allow a small degree of uncertainty in identification, as suggested by Lukacs and Burnham, 2005 (around 1-5%), as it is now possible to model this error rate (Link et al., 2010; Lukacs and Burnham, 2005; Wright et al., 2009; Yoshizaki et al., 2011). If the cost of adding a parameter (the error rate) is offset by the additional number of samples that can be retained, then the trade-off is of interest.

Of the approaches that incorporate the misidentification process into the analysis, Yoshizaki’s model (Yoshizaki et al., 2011) is the simplest, at the cost of discarding some data. It consists of removing all single-capture histories, allowing a more straightforward treatment of the uncertainty in the remaining dataset. But this model appears of little use as the amount of data that would be discarded often remains large. For example, Woodruff et al., 2016 collected pronghorns’ faeces over seven occasions, and approximately 30% of the identified individuals were detected only once. Thus, instead of deleting data, we favour modelling misidentifications in order to retain as much data as possible.

Wright’s model (Wright et al., 2009) uses the full dataset, but requires genotype replicates to estimate an error rate. This approach increases the costs, especially for large populations for which many samples may be obtained. The Latent Multinomial Model (LMM, Link et al., 2010) is a malleable framework that may be used to address the issue of keeping full data without additional costs. It estimates, in a Bayesian framework, the misidentification rate from the excess of single-capture histories compared to what is expected in the absence of misidentifications. It has recently received attention and has been extended by other publications to improve the computational efficiency of the model (Bonner et al., 2015; Schofield and Bonner, 2015) and to estimate population size in presence of individual heterogeneity (McClintock et al., 2014). However, one of the main hypotheses of the LMM is that individuals may only be seen once per occasion. In studies where DNA is extracted from faeces, the individuals can actually be ‘captured’ multiple times at the same occasion. As the position of misidentifications in capture histories remains unknown, it is not possible to keep only one observation per individual per capture session. The misidentifications will generate additional presence data and will ultimately lead to an overestimation of population size by the LMM.

In this paper, we develop an extension of the LMM to account for multiple observations of an individual on the same occasion. To illustrate the problem and our solution, we take the example of a study of the Eurasian otter (*Lutra lutra*) in Upper Lusatia, Saxony, Germany (Lampa et al., 2015). We first present the otter dataset as well as simulated data that cover more general situations. We then present the original model from Link et al., 2010, our extension and compare the results obtained from these models.

## Material and methods

### Otter data

Otters are nocturnal and elusive and pose challenges for (live-)trapping (Kruuk, 2006). Otter faeces (spraints) on the other hand are particularly conspicuous as they are used for intraspecific communication. Otters produce up to 30 spraints a day, tending to defecate on frequently visited visible terrestrial sites at specific locations throughout their home range. Thus, these spraints can be easily sampled for DNA-identification of the individuals. Otters are solitary and territorial, with home ranges that may overlap only marginally. Lampa et al., 2015 have collected spraints over five consecutive days (number of occasions: *T* = 5) from 2006 to 2012 (excluding 2009). Sampling was conducted in March (2006, 2010, 2011, 2012), April (2007) and May (2008). Each year, all ponds in the study area that contained water (covering 261–449 ha, depending on annual water management) were included in the sampling. During the five sampling days, all freshly deposited spraints around the ponds were collected. The datasets from each year were considered independently so the population can be considered closed (no birth, death, immigration, emigration) over T=5 five days. The authors considered it unlikely that repeated PCR could completely eliminate all genotyping errors due to relatively high genotyping error rates and low genotyping success rates. Thus, they used the error-incorporating misidentification model proposed by Lukacs and Burnham, 2005 (hereafter, model L&B), implemented in the MARK program (White and Burnham, 1999). However, this model does not adequately address capture histories resulting from misidentification as it ignores the dependence between the pair of histories created whenever an individual is misidentified (Link et al., 2010; Yoshizaki et al., 2011).

### Closed population capture recapture: model M_t_

When estimating population size *N* in a closed capture-recapture experiment (i.e. the population is assumed not to change) with the model known as *M*_*t*_ (Darroch, 1958; Otis et al., 1978), individuals are assumed to be observed (‘captured’) with probability *p*_*t*_ at occasion *t* for *t* = 1, 2, …, *T* and identified individually through tags/tracking devices or natural markings. Capture events are assumed independent between individuals and over time.

For each occasion, an individual is assigned 0 if it was not captured, or 1 if it was. This leads to 2^*T*^ possible distinct histories, including the unobservable all-zero history. They are represented by the sequence ***ω***_*i*_ = (*ω*_*i*,1_, …, *ω*_*i,T*_). Here, *y*_*i*_ is the number of individuals with history *ω*_*i*_ and 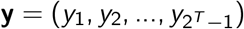. **y** follows a multinomial distribution with index *N* and cell probability

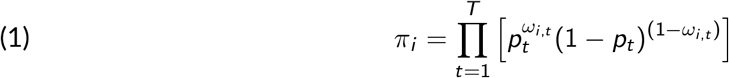

### Modelling misidentifications: model M_t,*α*_

To account for individual misidentifications, Yoshizaki et al., 2011 proposed the model *M*_*t,α*_, in which captured individuals are correctly identified with probability *α*. Misidentifications are assumed to create a new individual (a ‘ghost’). An individual cannot be mistaken as another, and two misidentifications cannot create the same ghost. Link et al., 2010 developed a latent structure for the model *M*_*t,α*_, which allows for Bayesian parameter estimation. In this structure, misidentifications are represented by 2 in the latent error histories ***ν***_*j*_ = (*ν*_*j*,1_, …, *ν*_*j,T*_). There are thus 3^*T*^ possible latent histories. The frequency of the latent error history *ν*_*j*_ is noted *x*_*j*_, and the vector of all latent error frequencies is 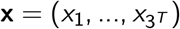. To make future developments of the model easier, the observation process of the likelihood is broken down into two parts, the capture process and the identification process. We followed Bonner et al., 2015 by introducing latent capture histories ***ξ***_*k*_ = (*ξ*_*k*,1_, …, *ξ*_*k,T*_). These are the true capture histories, i.e. in the absence of individual misidentifications, composed of 0 and 1. The frequency of the latent capture history *ξ*_*k*_ is noted *z*_*k*_, and the vector of all latent capture frequencies is 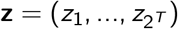.

In this model framework, the observed frequencies vector **y** is a known linear transformation **y** = **Ax** of the latent error frequencies **x** for a given matrix **A**. The constraint matrix **A** is (2^*T*^ − 1) × 3^*T*^ with a 1 in row *i* and column *j* if the latent error history *j* gives rise to the observed history *i*. All the other entries are zeros. The latent capture frequencies vector **z** is another linear transformation **z** = **Bx** for a given matrix **B. B** is 2^*T*^ × 3^*T*^ with 1 at row *k* and column *j* if the latent capture history *ξ*_*k*_ and the latent error history *ν*_*j*_ have the same capture pattern.

The joint distribution of **y, x** and **z** is

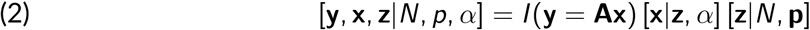

The probability of the capture process [**z**|*N*, **p**] is the same as in the closed CR model *M*_*t*_, using histories *ξ* and frequencies **z**. The capture likelihood is the following multinomial product where *π*_*k*_ is computed as in Equation 1, using history *ξ* instead of *ω*:

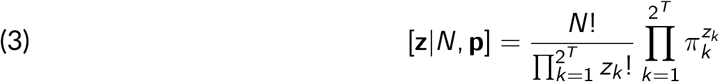

Bonner et al., 2015 gives the likelihood of the identification process, knowing the true captures:

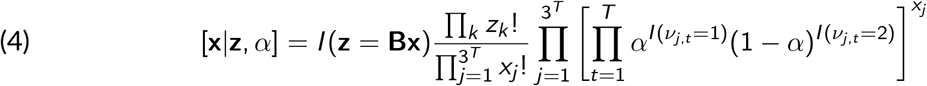

where *I* (*test*) is 1 if *test* is true, and is 0 otherwise.

The ratio of factorials accounts for the relabelling of the marked individuals that would produce the same counts in **x** and **z**.

The full likelihood is obtained by summing the conditional [**y**|**x, z**, *N, p, α*] over all values of **x** belonging to the set ℱ_**y**_ = {**x**|**y** = **Ax**}:

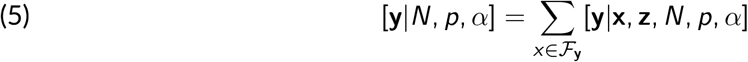

Note that there is no summation over **z** in this equation because **z** is the result of a deterministic function of **x** (**z** = **Bx**).

The feasible set ℱ_**y**_ is complicated to enumerate, which makes the likelihood (Equation 5) almost intractable in terms of computation. Maximum likelihood estimation is therefore not practical. Conveniently, Link et al., 2010, Schofield and Bonner, 2015 and Bonner et al., 2015 show how a Markov chain Monte Carlo (MCMC) can be used in this context. The Markov chain allows for the estimation of the posterior density:

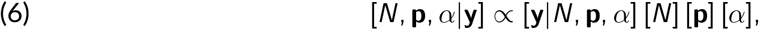

where [*N*], [**p**] and [*α*] denote the priors on population size, capture probability and identification probability.

The complete algorithm is detailed in Section A of the Supplementary Material.

### Extension to repeated observation: model *M*_*λ,α*_

In this section, we consider the case in which several captures of the same individual can occur at the same occasion. In this case, the observable history *ω* of the individual is composed of counts representing the number of times an individual was observed and identified at each occasion. For example, we might observe the following individual history: (0, 2, 0, 3, 1), where 2 is the number of times the individual is detected at occasion 2. In a monitoring in which individuals can be ‘captured’ several times on one occasion, one of the captures might result in a misidentification and the creation of a ghost history, while the other captures are correct identifications. In contrast to the model *M*_*t,α*_, an individual producing a ghost at a given occasion can be seen at this occasion if another capture of it results in a correct identification. Due to the new structure of the data, we must modify the notations used for misidentifications: we cannot use ‘2’ for indicating a misidentification. In addition, a history could contain several misidentifications for the same occasion. To represent latent error histories *ν*_*j*_, we note the total number of observations of an individual on each occasion with the number of these observations that resulted in misidentification as a superscript. For example, the previous observed history might have been generated by the latent error history (1^(1)^, 2, 0, 3, 3^(2)^). In this example, the observation at the first occasion and two observations at the fifth occasion were misidentified, resulting in zero observations at occasion 1 and one observation at occasion 5. The latent capture history *ξ*_*k*_ is the same as the latent error history without the superscripts. In our example, the latent capture history is (1, 2, 0, 3, 3).

In this section, we make the same assumptions about the misidentifications as in the model *M*_*t,α*_. Identifications are independent, misidentifications always result in ghosts (i.e. false individuals), and ghosts cannot be seen again (i.e. ghosts have exactly one observation in their history). As in the model *M*_*t,α*_, the observed histories can be summarized in the frequency vector **y**.

The different latent error histories possibly responsible for the observed histories can be summarized in the frequency vector **x**. As in the model *M*_*t,α*_, the vector **y** is the linear transformation of the vector **x**: **y** = **Ax**, where **A** is a known matrix. Table 1 shows an example of **y, x** and **A** in the case where the three histories (1, 2), (0, 3) and (1, 0) were observed, and there is only one history (1, 0). In this case, there is only one possible ghost, but we do not know which history might have generated it.

**Table 1.**
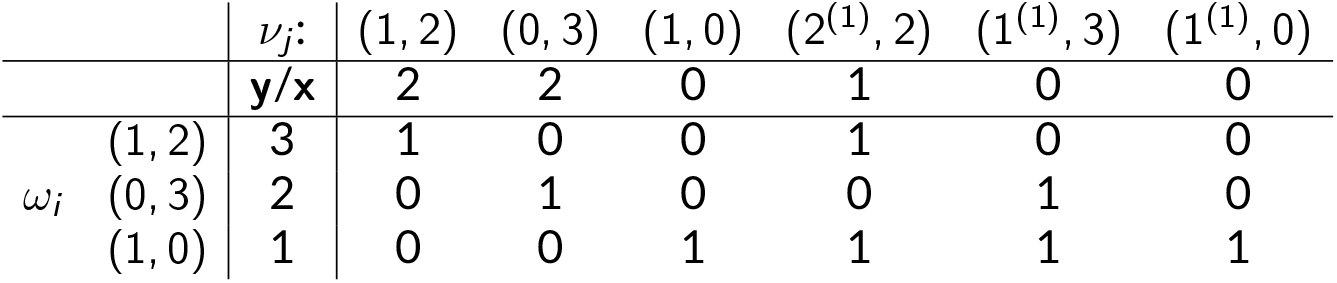
Example of a correspondence between y and x: the first and second rows are, respectively, the possible latent error histories and a possible latent error history frequencies. The first and second columns are, respectively, the observed histories and the observed history frequencies. The rest of the table is the matrix **A**.

Similarly, the latent capture histories can be summarized in the frequency vector **z**. Also, **z** is a linear transformation of **x**: **z** = **Bx**. Table 2 illustrates the example from above, showing possible **z** and **B**.

**Table 2.** Example of a correspondence between x and z: the first and second rows are, respectively, the latent error histories and a possible latent error history frequency. The first and second columns are, respectively, the latent capture histories and the latent capture history frequencies corresponding to the given latent error history frequencies. The rest of the table is the matrix **B**.

| | | $\nu_j:$ | (1, 2) | (0, 3) | (1, 0) | $(2^{(1)}, 2)$ | $(1^{(1)}, 3)$ | $(1^{(1)}, 0)$ |
| --- | --- | --- | --- | --- | --- | --- | --- | --- |
| | | $\mathbf{z}/\mathbf{x}$ | 2 | 2 | 0 | 1 | 0 | 0 |
| $\xi_k$ | (1, 2) | 2 | 1 | 0 | 0 | 0 | 0 | 0 |
|  | (0, 3) | 2 | 0 | 1 | 0 | 0 | 0 | 0 |
|  | (1, 0) | 0 | 0 | 0 | 1 | 0 | 0 | 1 |
|  | (2, 2) | 1 | 0 | 0 | 0 | 1 | 0 | 0 |
|  | (1, 3) | 0 | 0 | 0 | 0 | 0 | 1 | 0 |

Here, ***λ*** = (*λ*_1_, …, *λ*_*T*_) is the set of parameters involved in the capture process, modelled by a Poisson process (each *λ*_*t*_ being the parameter of the Poisson process at occasion *t*). The joint distribution of **y, x** and **z**, is given by the Equation 7.

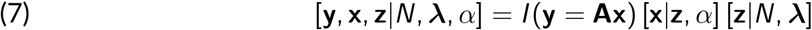

We now describe the two elements that change compared to model *M*_*t,α*_, i.e. [**z**|*N*, ***λ***] and [**x**|**z**, *α*].

- [**z**|*N*, ***λ***]

We model the true number of observations of an individual at an occasion with a Poisson distribution. The probability *π*_*k*_ that an individual has a given latent capture history *ξ*_*k*_ is given by:

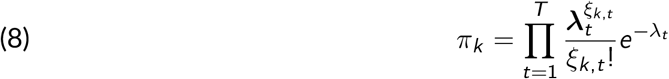

The capture likelihood has a multinomial form:

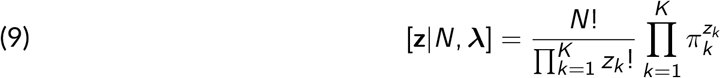

where *K* is the number of histories *ξ*_*k*_ with positive count. Despite being an infinite number of possible latent capture history *ξ*_*k*_, the product terms become 1 when *z*_*k*_ = 0 and thus need not be computed.

- [**x**|**z**, *α*]

All actual captures are potentially subject to misidentifications. If *o*_*j,t*_ is the number of correct identifications for individuals with history *ν*_*j*_ at occasion *t*, then, knowing the true number of captures, the probability that it was correctly identified *o*_*j,t*_ times is binomial. Thus:

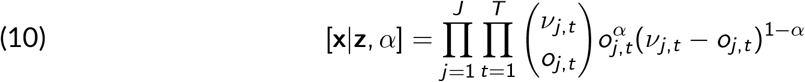

where *J* is the number of histories *ν*_*j*_ with positive count.

In addition to this model, we also adapted the algorithm to estimate the parameters. The full algorithm is presented in Section B of the Supplementary Material.

### Simulations

To validate our approach for a broader range of situations than the otter case, we conducted a simulation study. We simulated datasets for a population of *N* = 500 individuals. The number of captures of each individual at each occasion (0 or more) were sampled from Poisson distributions with time-dependent parameters *λ*_*t*_. We aimed for four different capture rates (with the meaning ‘seen at least once’ or ‘not seen at all’ on one occasion) of 0.1, 0.2, 0.3 and 0.4, so we set *λ*_*t*_ = 0.11, 0.23, 0.36, 0.51 respectively (so on average, with *λ*_*t*_ = 0.11, 10% of the individuals will be seen at least once and 90% will remain unseen). Three different simulated identification rates were used with values *α* = 0.8, 0.9, 0.95. We simulated 10 datasets for these 12 scenarios.

Population size is not expected to influence parameter identifiability or bias, provided that sampling effort and detection probabilities remain comparable (Otis et al., 1978). We therefore did not vary the population size as our primary objective was to evaluate the model’s ability to produce unbiased estimates under different levels of sampling intensity and misidentification, rather than to assess its sensitivity to population size itself. However, to facilitate comparison with the otter dataset, which involves a relatively small population, we also conducted additional simulations with a population size of 50 individuals. This analysis is presented in the Supplementary Material.

### Implementation and MCMC configuration

We used NIMBLE (Valpine et al., 2017) to implement the models. The advantage of NIMBLE is that it allows writing all the samplers of the MCMC (mandatory here) and new distributions. Thus, we wrote the likelihood of the model and the sampler of **x**. We were also able to rewrite all the Gibbs samplers of the MCMC for maximum computational efficiency. To improve efficiency, all observable histories with a count of zero were ignored, i.e. their corresponding rows and columns in the matrices **A** and **B** were deleted, as suggested in Schofield and Bonner, 2015.

All priors were taken to be uninformative or vague. The *p*_*t*_ and *α* parameters had a *β*(1, 1) distribution prior and the *λ*_*t*_ parameters had a *γ*(1, 1) distribution prior. For the simulated datasets, the *M*_*λ,α*_ model was run over 1E6 iterations after a burn-in period of 100,000 iterations. To simplify the computation of statistics and use less storage, the chains were thinned by a factor of 1/100. Yoshizaki’s model was run over 10,000 iterations after a burn-in period of 1000 iterations. The *M*_*t*_ model was run over 100,000 iterations after a burn-in period of 10,000 iterations, and the chains were thinned by a factor of 1/10.

For the otter dataset, the *M*_*λ,α*_ model, Yoshizaki’s model and the *M*_*t*_ model were run over 10,000 iterations after a burn-in period of 1,000 iterations.

For the otter dataset, we ran three chains with different starting values. In contrast, for the simulations - where the true values are known and previous tests showed that two chains were sufficient to assess convergence - we used only two chains. The first starting point had **x** initialized as the set of observed histories, as if there were no errors. In the second starting point (and third for the otter dataset), using the algorithm that allows moving along the feasible set F_**y**_ (Supplementary material, section B, step 5) we arbitrarily added 40 random errors for the simulations and 5 for the otter (unlike in the MCMC, we systematically accepted the proposals of the algorithm). This procedure preserved the structure of the observed data.

## Results

### Simulations results

We evaluated our extension of the LMM, *M*_*λ,α*_, against the model proposed by Yoshizaki et al., 2011, in which all histories with a single capture are excluded. We also tested the standard capture-recapture model *M*_*t*_ (that ignores misidentifications) to demonstrate the importance of accounting for misidentifications.

Running two chains of 1,100,000 iterations for the model *M*_*λ,α*_ on a 3.0GHz Intel processor took less than 10 minutes. We checked the convergence graphically by looking at the *N* chains, as the estimated number of individuals *N* is the slowest moving parameter. We also checked whether the 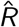 of the *N* chains was under 1.1. As the *N* chains are highly correlated, we checked whether the resulting effective sample size was at least over 200. More iterations could be used to increase sample size but running them on simulations would have taken much more time for very little additional precision. These “low effective sample size” simulations show situations where the MCMC takes too many iterations to fit the model, which could indicate a lack of data. The model *M*_*t*_ always converged perfectly (except for two simulations with five occasions and *λ* = 0.11) and there was no evidence of any problems. Yoshizaki’s model converged for all simulations except for some with 5 or 7 occasions, *α* = 0.8 and *λ* = 0.11. But some *N* chains for *λ* = 0.11 show peaks above ten times the mean of the estimates (one chain has a peak value over 20.000 while the median is around 200). For the *M*_*λ,α*_, the MCMC also converged for almost all simulations except again for some with 5 or 7 occasions, *α* = 0.8 and *λ* = 0.11. The convergence was always faster for higher identification rates.

The population size estimates for all models are shown in Figure 1. The model *M*_*t*_ always overestimates the population size, up to a factor of two. Lower identification rates and higher capture rates both lead to greater overestimation. Yoshizaki’s model and the *M*_*λ,α*_ model have the same trend of estimates: for very low capture rates, the population size estimates are highly biased, down to less than half the true population size. For a capture rate of 0.2, there is only a small bias with seven or fewer occasions, and no bias with nine occasions. There was no bias for higher capture rates.

**Figure 1.**
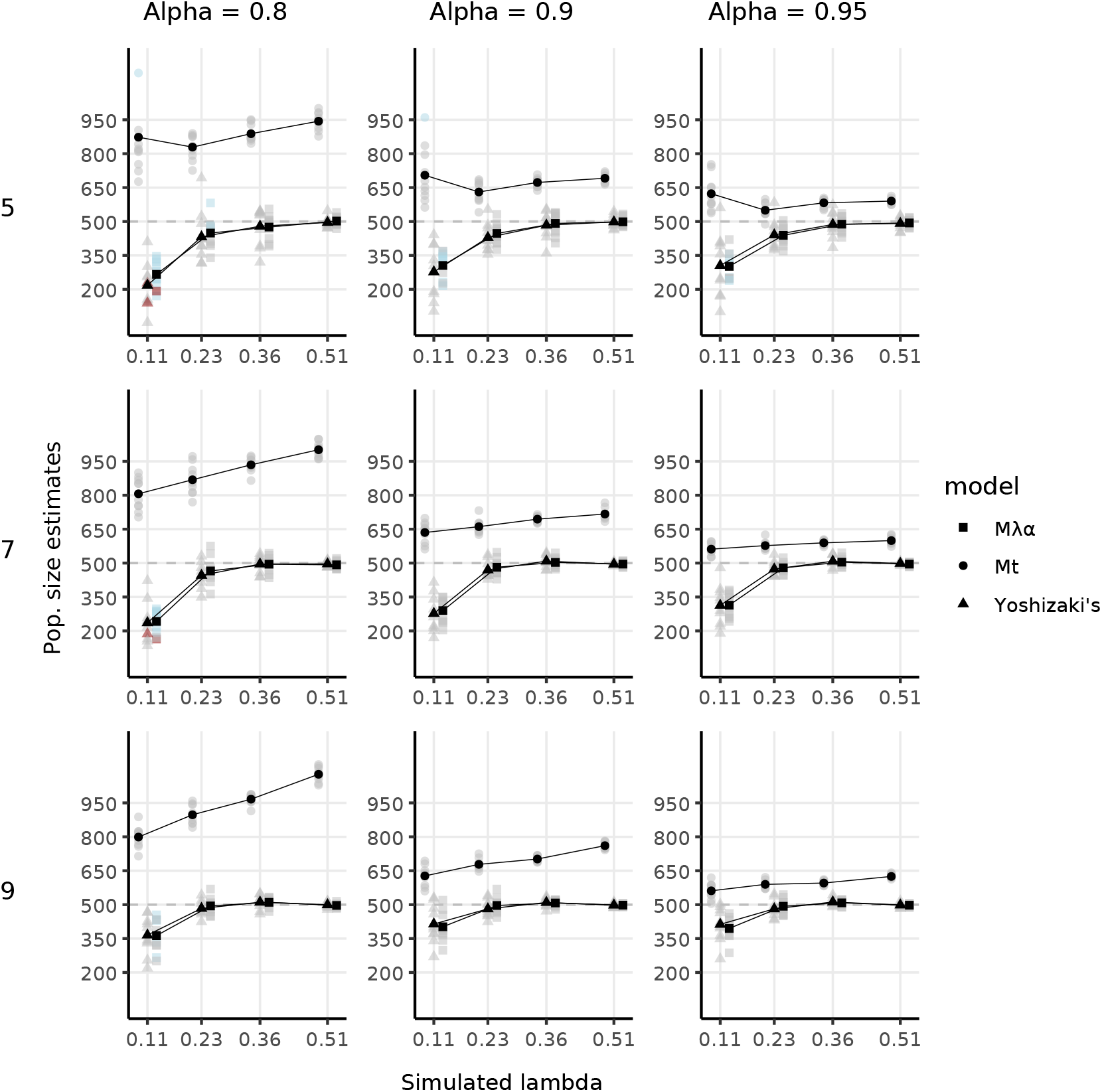
Study of performance of the population size estimators of models *M*_*λ,α*_, *M*_*t*_, and Yoshizaki’s using simulations (see main text for details) (y axis) under different levels of capture probability (x axis), identification probability (columns), and number of capture occasions (lines). Horizontal dashed lines indicate true population size. Grey, red and blue symbols show simulation-specific estimates of the population-size posterior mean. Red symbols indicate that the N chains had 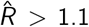. Blue symbols indicate 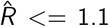 but an effective sample size under 200. Black symbols connected by lines show the mean estimates averaged across simulations.

The credible intervals estimated by the model *M*_*λ,α*_ are smaller than that estimated by Yoshiza-ki’s model for most scenarios with seven or fewer occasions. They are slightly larger with nine occasions, but the true population size is more often in the credible interval estimated by the model *M*_*λ,α*_ as shown in Table 3.

**Table 3.** Percentage of simulations for which the true population size was inside the estimated credible interval. See the supplementary material section C for a detailed table depending on all parameters values.

| S | $M_{\lambda, \alpha}$ | Yoshizaki | $M_t$ |
| --- | --- | --- | --- |
| 5 | 87.5 | 86.7 | 13.3 |
| 7 | 84.2 | 84.2 | 7.5 |
| 9 | 95.0 | 77.5 | 5.0 |

### Otter results

The estimates are shown on Figure 2. For 2011, all models estimate a similar population size. For all other years, the *M*_*t*_ model produces higher estimates of the population size compared to the other models. For most years, the *M*_*λ,α*_ model and Yohizaki’s model have similar estimates. The minimum population size (numbers of observed individuals that cannot be ghosts considering the hypothesis of the LMM) are 16, 22, 14, 16, 23, 18 for each year respectively. For 2006, 2007 and 2010, the population size estimated by Yoshizaki’s model is yet below this minimum.

**Figure 2.**
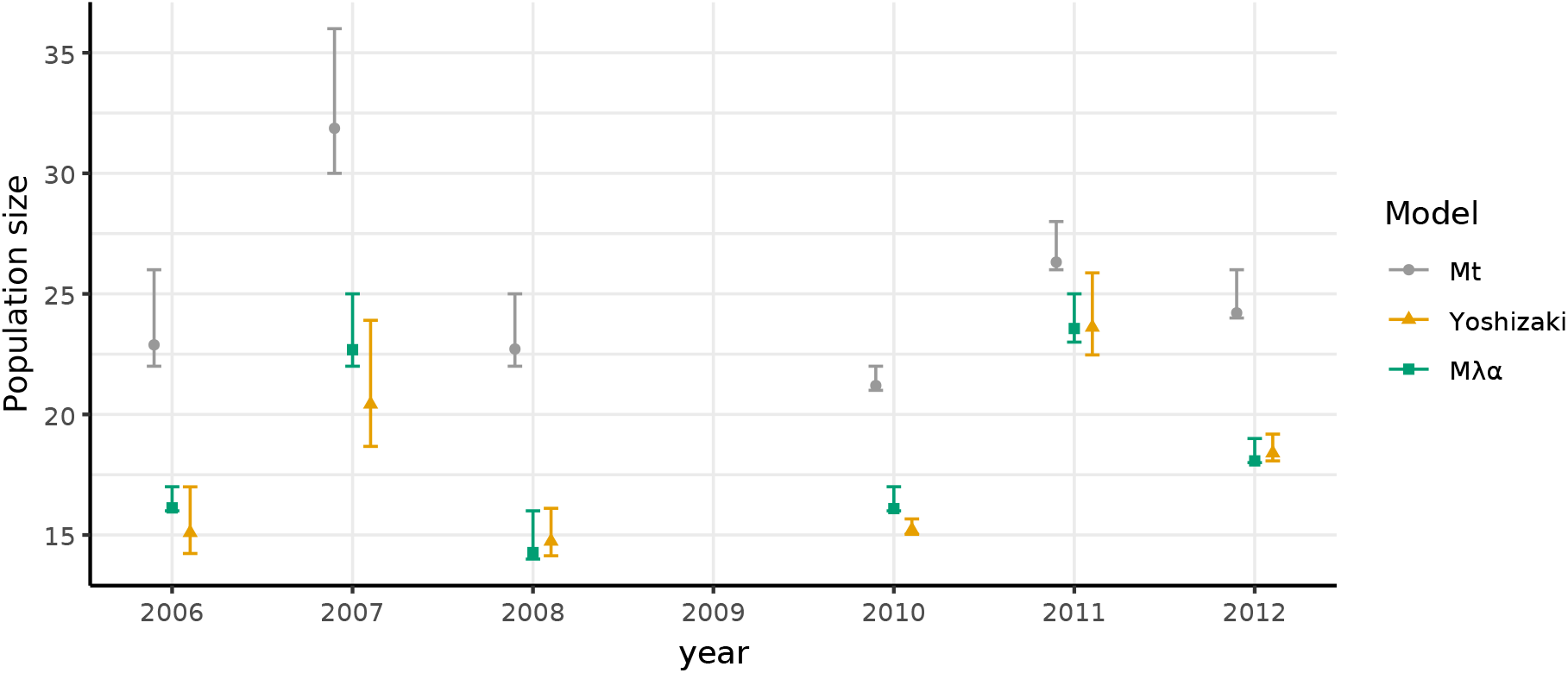
Otter population size estimations. The error bar are the 95% confidence intervals.

The capture parameters estimated by models are shown on Figure 3 and given in the supplementary material section D. For all years, the confidence intervals produced by Yoshizaki’s model overlap those produced by the other models. For 5 different occasions (on different years), the confidence intervals produced by the models *M*_*t*_ and *M*_*λ,α*_ do not overlap (and for three more occasions the overlap is less than 5% of the interval length). For almost all occasions and all years, the *M*_*t*_ model gives the lowest capture rate mean estimates, ranging from 0.4 to 0.7. The *M*_*λ,α*_ model gives much higher capture rates, between 0.6 and 0.9, except for one occasion at 0.4. The estimated *λ*_*t*_ range from 0.5 to 2.4. Yoshizaki’s model often gives estimates between the other two models, ranging from 0.5 to 0.9. About overall detection, under the misidentification hypotheses, Yoshizaki’s model and the *M*_*λ,α*_ model both estimate that all individuals were seen at least once. The *M*_*t*_ model also estimates that all individuals were seen at least once.

**Figure 3.**
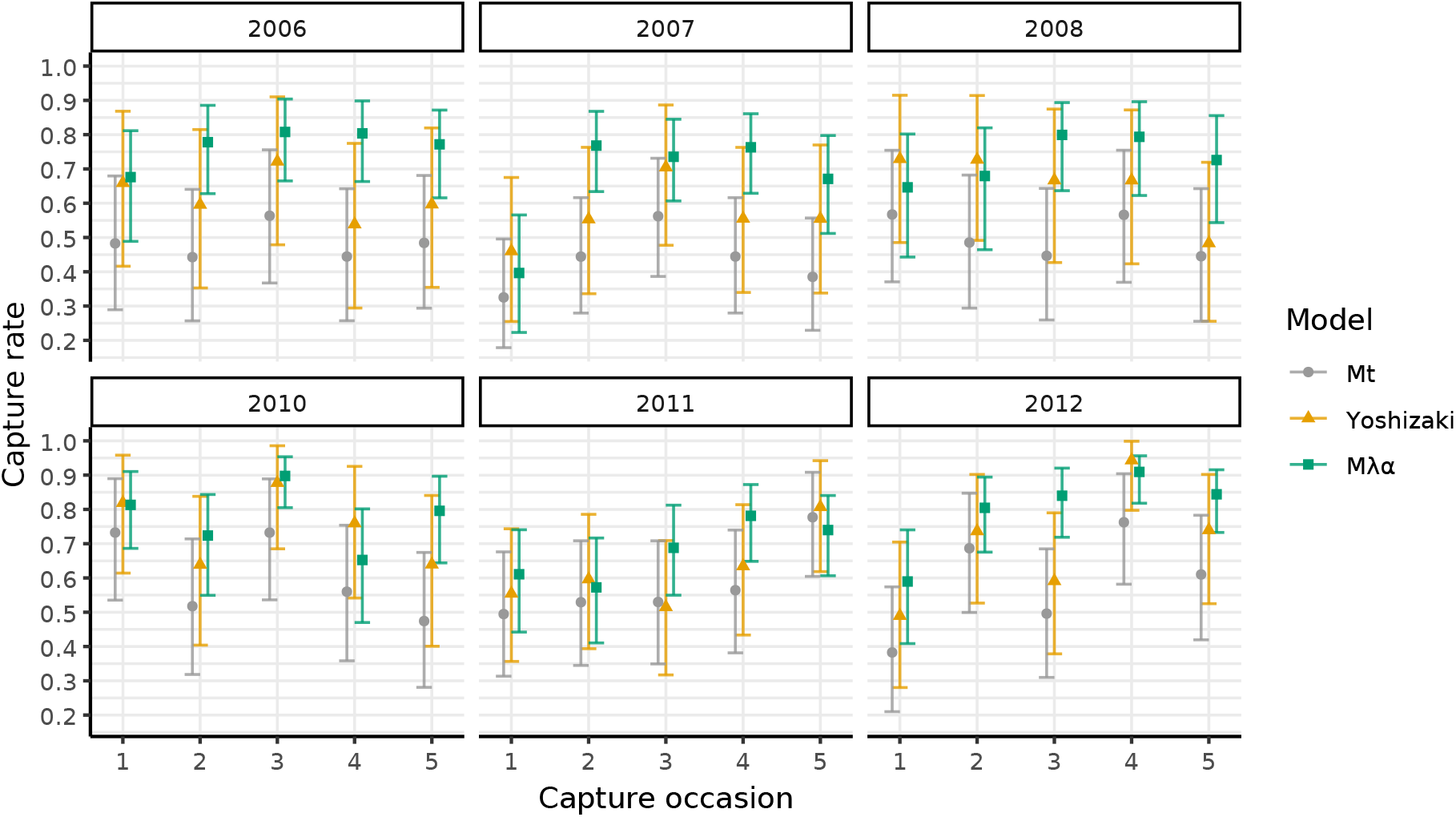
Capture rate estimates for the otter dataset. The error bar are the 95% confidence intervals. The values given for the *M*_*λ,α*_ model are the computed probability of having “captured” an individual at least once at an occasion instead of the mean number of capture so that it can be compared with the other models.

Under the misidentification assumptions of the LMM, the maximum possible numbers of misidentifications were 6, 8, 8, 5, 3, and 6 for each year, respectively, corresponding to the counts of single-capture histories. The respective maximum error rates are 11%, 12%, 14%, 8%, 4% and 8%. The number of estimated misidentifications is shown in Table 4. For all years, the model *M*_*λ,α*_ estimated that all the histories with a single capture were probably ghosts. In contrast, for most years, the mean estimation of identification rates from model L&B - that inadequately accounts for misidentification under our assumptions - indicated more misidentifications than the theoretical maximum. For this to be the case, misidentifications would have to occur in histories with more than one capture, thereby violating either the assumption that individuals cannot be confounded or the assumption that misidentifications generate distinct ghost histories.

**Table 4.**
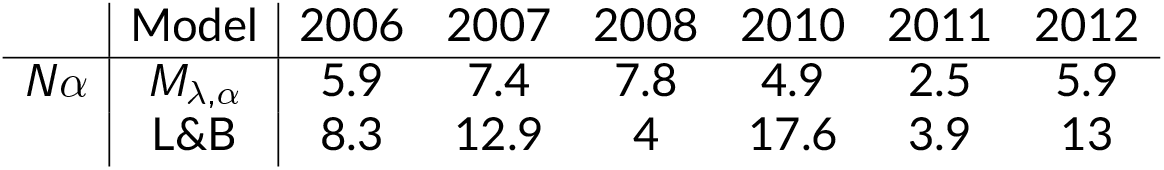
Number of misidentifications estimated. For the *M*_*λ,α*_ model it’s the mean number, for the L&B it’s the estimated misidentification rate multiplied by the number of captures.

## Discussion

In this paper, we treat the ecological question of population size estimation with capturerecapture techniques when the individuals are identified using DNA extracted from non-invasive samples such as faeces, hair, or feathers. We develop an extension of the LMM (Link et al., 2010) to account for repeated observations on a given sampling occasion. We apply our model to an otter dataset and conduct a simulation study to demonstrate its efficiency and usefulness.

Concerning the otter dataset, all years except maybe 2011 most probably contained misidentifications according to Yoshizaki’s and our *M*_*λ,α*_ model. Ignoring the misidentifications (*M*_*t*_ model) leads to an estimate up to 30% higher than when accounting for misidentifications. This example, and the overestimation of the population size in the simulations, should be sufficient to induce researchers to account for potential misidentifications.

Yoshizaki’s model relies on deleting unsure data but it also leads to delete sure and available data. For example, histories of individuals seen at only one occasion but several times (like “03000”) cannot be ghosts. However, these histories are still removed from the data. All additional captures, are also removed. For example, the history “20130” is simplified into “10110”. This loss of information can lead to an underestimation of the population size, as happened with the 2006, 2007 and 2010 otter datasets. It might also be responsible for higher uncertainty as observed in the simulation study. In contrast, our model can make use of all the available information to get better estimates of population size.

The otter data set presents features that have been identified as favouring a good performance of our model: the capture rates were estimated over 0.5, the highest simulated value, and the identification rate was around the highest simulated (95%). However, the population size was very small compared to the simulations. A small simulation study close to the otter dataset with the lowest capture rate estimated for it (*N* = 30, *λ* = 0.5, *α* = 0.95), presented in the supplementary material section E, shows that a small population size can induce a small bias (average -6%). However the bias is not systematic and for most simulations (47/50), the true population size lies in the credible interval.

While being the first model to allow simultaneous samples handling (i.e. repeated sampling on the same “capture” occasion), our model currently lacks certain features present in previous models. For instance, it assumes capture homogeneity among individuals. However, capture heterogeneity is not uncommon: for the years 2007 and 2008, Lampa et al., 2015 identified the *M*_*h*_ model that considers capture heterogeneity between individuals as the best. This could be due to some spatial distribution of the individuals which gives better chances to collect samples from individuals more present along the transects used to collect data. The years 2007 and 2008 correspond to the highest recorded water surface areas, which may indicate increased hydrological connectivity, particularly with ponds outside the study area. This could partly explain the observed heterogeneity, as individuals located near the boundaries of the monitored area may have moved more frequently beyond its limits, resulting in lower capture probabilities. McClintock et al., 2014 showed how to model capture heterogeneity for the *M*_*t,α*_ model. What they did could be extended to our *M*_*λ,α*_ model.

We move now to discussing the main hypothesis we made in modelling the collection of several samples from the same individual at the same occasion. Indeed in using a Poisson process, we implicitly assumed independence of the samples. However, there could be significant over or under dispersion in the count data. A few simulations with 50 or 100 individuals and the variance up to three times the mean (for the capture counts) showed no bias in the estimates and showed the robustness of the method in presence of non-independent samples. Moreover, if needed the Poisson distribution could be changed to another distribution that accounts for overdispersion like the negative binomial.

For the identification process, we assume that misidentifications cannot be repeated. This implies that the identification panel (either using microsatellites or SNPs) should be strong enough to allow a good distinction between individuals (and close kins if necessary). The minimal number of loci to distinguish individuals should be fixed so that the probability of making the same mistake twice is very low, ensuring misidentifications are not repeated. And by keeping a threshold low enough for the Probability of Identity (Taberlet and Luikart, 1999), we can ensure that no two individuals are confused. To reduce the risk of misidentification, the authors of the otter study first discarded low-quality samples based on predefined thresholds (Lampa et al., 2013). They then applied a screening protocol with five amplification steps, repeating any sample with at most two missing loci after the fifth step until a reliable genotype was obtained. Because only samples with complete genotypes were retained, the likelihood of confusing individuals is very low. For the same reason, the probability of reproducing the same genotyping error across several samples from the same individual - and thus repeating the same misidentification - is expected to be negligible.

If misidentifications were significantly repeated, the model would most likely overestimate population size since it would consider histories like “020” - where the “2” results from two identical misidentifications - as a real separate individual. However, the model could be extended to consider histories with more than a unique capture as potential ghosts, but the power to detect this event would probably be extremely low. If individuals were to be confused, the *M*_*t,α*_ or *M*_*λ,α*_ models would not account for it, probably biasing the estimates. The model from Bonner et al., 2015 that deals with that kind of misidentification could be extended to repeated observations on the same occasion as we did here if necessary.

Finally, the question addressed in this paper is that of population size estimation. However, many studies focus on estimating survival rather than population size and face the same problem of potential misidentifications. Therefore, it seems interesting and important to develop an extension of the LMM that would estimate survival with repeated observations.

To conclude, we show that our *M*_*λ,α*_ model is a great tool to analyse data obtained using DNA from samples such as faeces, hair or feathers and to deal with the misidentification problem while using all the data available. Moreover, using such a model can bring great benefits, from simply confirming the absence of misidentification to accounting of this issue when analysing the data - thus using misidentification-prone data that would have been discarded.

## Acknowledgements

We would like to thank Daniel B. Turek at Williams College Department of Mathematics and Statistics, without whom the NIMBLE implementation would have taken much more time. We would also like to thank the editor and the two reviewers that helped us a lot with improving the paper. Finally, we thank all the administration, engineers, technicians and cleaning people that kept the lab running while we were working on this.

Preprint version 4 of this article has been peer-reviewed and recommended by Peer Community In Ecology (https://doi.org/10.24072/pci.ecology.100744).

## Data, script, code, and supplementary information availability

Otter data from Lampa et al., 2015 are available online (https://doi.org/10.1371/journal.pone.0125684.s002)

Script and codes are available online (https://doi.org/10.5281/zenodo.15575907)

Supplementary information is available online (https://doi.org/10.1101/2024.06.12.598605 ; https://www.biorxiv.org/content/10.1101/2024.06.12.598605v2.supplementary-material)

