## Supplementary material for "Population size estimation when multiple samples carrying the risk of misidentification are taken within the same capture occasion from the same individual"

### Estimating population size with repeated observations of non-invasive sampling in presence of misidentification

1: CEFE, Univ Montpellier, CNRS, EPHE, IRD – Montpellier, France

#### Supplement A: Bayesian Parameter Estimation for $M_{t,\alpha}$

The MCMC is constructed this way:

(1) Let $\beta(a_0^t, b_0^t)$ denote the beta prior on $p_t$ and $\beta(a_0^\alpha, b_0^\alpha)$ denote the beta prior on $\alpha$.

(2) Initialize all parameters as well as a set of latent histories satisfying **y**=**Ax**. Such a set can be obtained by assuming that no errors have been made. The latent frequencies of the histories containing 2's are 0 and all others match the observed frequencies one to one. To run multiple chains with different initializations, one can take the previous initialization of **x** and follow the later steps of (5) only adding misidentifications to the set and always accepting the proposed ones without going through the Metropolis-Hasting acceptance. In the initial latent set, fix the number of unseen individuals to a realistic random number.

(3) Sample the capture rate using Gibbs sampling as shown in Link et al., 2010. The likelihood being multinomial, it follows that the beta priors lead to full conditional beta posterior distribution:

$$p_t|\mathbf{z} \sim \beta(a_0^t + a^t, b_0^t + b^t)$$

where $a^t$ is the number of captured individuals at time $t$ and $b^t$ the number of unseen individuals at time $t$ (including those never seen). $a^t = \sum_n I(\xi_{n,t} = 1)$ and $b^t = \sum_n I(\xi_{n,t} = 0)$.

(4) Sample the identification rate using Gibbs sampling. Similar to the capture rate, it has a full conditional beta posterior distribution:

$$\alpha|\mathbf{x} \sim \beta(a_0^\alpha + a^\alpha, b_0^\alpha + b^\alpha)$$

where $a^\alpha = \sum_j x_j I(\nu_{j,t} = 1)$ is the total number of correct identifications and $b^\alpha = \sum_j x_j I(\nu_{j,t} = 2)$ the total number of misidentifications.

(5) Sample jointly $N$ and **x** since the number of errors in **x** changes the population size. Sampling **x** requires being able to sample from $\mathcal{F}_\mathbf{y}$. Link et al., 2010 suggests sampling moves from the null space of the matrix **A**, $Ker_\mathbb{Z}(\mathbf{A})$ (*i.e.* from the set of **x** such as $\mathbf{Ax} = 0$),

$$Ker_\mathbb{Z}(\mathbf{A}) = Ker(\mathbf{A}) \cap \mathbb{Z}^d = \{\mathbf{x} \in \mathbb{Z}^d | \mathbf{Ax} = 0\}$$

and adding or subtracting them to the current **x** in the MCMC. Schofield and Bonner, 2015 showed that if the basis of $Ker_\mathbb{Z}(\mathbf{A})$ is not carefully chosen, some parts of the space $\mathcal{F}_\mathbf{y}$ could be disconnected from the others and the Markov chain would only explore subspaces, depending on the initial **x**, possibly leading to biased estimates. They proposed to sample moves from the Markov basis of **A** Diaconis and Sturmfels, 1998, a set in $Ker_\mathbb{Z}(A)$ that connects all $\mathcal{F}_\mathbf{y}$ irrespective of the values in **y**. Such a basis ensures that the whole set $\mathcal{F}_\mathbf{y}$ is connected by single moves and that no move can leave the set. The disadvantage is that the computation of this Markov basis is heavy and algebraic software like 4ti2 team, n.d. will not be able to compute it for $T \geq 5$.

Bonner et al., 2015 proposed a mechanism to avoid computing this basis. It consists in sampling from the dynamic Markov basis Dobra, 2012 which is the set of moves $M(x)$ connecting each **x** to some neighbors. The algorithm randomly adds or removes an error from the set of latent histories. To add an error, the authors sample a history that might have generated a ghost (*i.e.* a history containing a 0), and "merge" it with a potential ghost (*i.e.* replace the 0 with a 2 and remove the ghost history). To remove an error, they sample a history containing a 2, replace it with a 0 and add a history with a unique capture (coded 1) at that time.

More formally, follow the steps:

(a) Define:
- $\nu_{1t}$ the history with a unique capture at time $t$ (potential ghost),
- $\chi_{0,t}(\mathbf{x}) = \{\nu|\nu_t = 0, x_\nu > 0, x_{\nu_{1t}} > 0\}$ the set of histories that *potentially* generated a ghost at occasion $t$, for the given $\mathbf{x}$,
- $\chi_{2,t}(\mathbf{x}) = \{\nu|\nu_t = 2, x_\nu > 0\}$ the set of histories *containing* a ghost at time $t$, for the given $\mathbf{x}$.

(b) With probability 0.5, go to (i), otherwise go to (ii).
  (i) Add a misidentification (i.e. a ghost) to the latent set.
  - Sample $\nu_0 \in \chi_{0.}(x) = \bigcup_t \chi_{0,t}(x)$.
  - Sample $t \in \{t|\nu_{0,t} = 0, x_{\nu_{1t}} > 0\}$.
  - Define $\nu_2 = \nu_0 + 2\nu_{1t}$.
  - Define the move $b_{\nu_0,\nu_1,\nu_2} = (-1, -1, +1)$.

  (ii) Remove a misidentification from the latent set.
  - Sample $\nu_2 \in \chi_{2.}(x) = \bigcup_t \chi_{2,t}(x)$.
  - Sample $t \in \{t|\nu_{2,t} = 2\}$.
  - Define $\nu_0 = \nu_2 - 2\nu_{1t}$.
  - Define the move $b_{\nu_0,\nu_1,\nu_2} = (+1, +1, -1)$.

(c) Define $\mathbf{x}' = \mathbf{x}^{(k-1)} + b$.

(d) Calculate $\mathbf{z}' = \mathbf{B}\mathbf{x}'$ and $N' = \sum \mathbf{x}'$.

(e) With probability $r_1$, set $\mathbf{x}^k = \mathbf{x}'$, $\mathbf{z}^k = \mathbf{z}'$ and $N(k) = N'$. Otherwise set $\mathbf{x}^k = \mathbf{x}^{k-1}$, $\mathbf{z}^k = \mathbf{z}^{k-1}$ and $N(k) = N^{(k-1)}$.

$$(1) \quad r_1 = min\left(1, \frac{[\mathbf{y}|\mathbf{x}', \mathbf{z}', N', p, \alpha]}{[\mathbf{y}|\mathbf{x}^{(k-1)}, \mathbf{z}^{(k-1)}, N, p, \alpha]} \frac{q(\mathbf{x}^{(k-1)}|\mathbf{x}')}{q(\mathbf{x}'|\mathbf{x}^{(k-1)})}\right)$$

The proposal densities are calculated by multiplying the probabilities of each sampling step used to define the move. They are, in order: the probability of adding (or removing) an error, the probability of choosing the $\nu_0$ (or $\nu_2$) and the probability of choosing the $t$ knowing the sampled $\nu$. When adding an error, the proposal density $q$ is:

$$(2) \quad q(x'|x^{k-1}) = \frac{0.5}{\#\chi_{0.}\#\{t|\nu_{0,t} = 0, x_{\nu_{1t}} > 0\}}$$

and when removing an error, is:

$$(3) \quad q(x'|x^{k-1}) = \frac{0.5}{\#\chi_{2.}\#\{t|\nu_{2,t} = 2\}}$$

where $\#S$ denotes the cardinality of $S$.

(6) Sample the number of unseen individuals $x_0$:
  (a) set $\mathbf{x}' = \mathbf{x}$, $\mathbf{z}' = \mathbf{z}$ and $x_0$ the number of unseen individual in $\mathbf{z}$ (and $\mathbf{x}$),
  (b) sample a move $c \in [-D, D]$ where D is fixed integer,
  (c) define $x_0' = x_0 + c$,
  (d) set the number of unseen individuals in $\mathbf{x}'$ and $\mathbf{z}'$ to $x_0'$,
  (e) accept it with probability $r_2$:

$$(4) \quad r_2 = min\left(1, \frac{[\mathbf{z}'|N', p]}{[\mathbf{z}|N, p]}\right).$$

(7) repeat steps 3 to 6 as much as needed.

#### Supplement B: Bayesian Parameter Estimation for $M_{\lambda,\alpha}$

From the algorithm used for model $M_{t,\alpha}$, we need to change how we sample $\boldsymbol{\lambda}$ which wasn't in the original model, but also the way we propose a new **x**. The way we sample $\alpha$ does not really change.

The MCMC is constructed this way:

(1) Let $\lambda_t \sim \Gamma(\alpha_0^{(t)}, \beta_0^{(t)})$ be the Gamma prior over $\boldsymbol{\lambda}$, and $\beta(a_0^\alpha, b_0^\alpha)$ denote the beta prior on $\alpha$.

(2) Initialize all parameters as well as a set of latent histories satisfying **y**=**Ax**. Such a set can be obtained by assuming that no mistakes were made. The latent frequencies of the histories containing 2's are 0 and all the other match the observed frequencies one-to-one. In order to run several chains with different initialisations, one can take the previous initialisation of **x** and follow the later steps of (5) by only adding misidentification to the set and always accepting the proposed ones without going though the Metropolis-Hasting acceptance. In the initial latent set, fix the number of unseen individual to a random realistic number.

(3) Sample the capture parameter $\boldsymbol{\lambda}$ with Gibbs sampling. The likelihood being a product of Poisson, it follows that the Gamma priors lead to full conditional Gamma posterior distribution:

$$\lambda_t|\mathbf{z} \sim \Gamma(\alpha_0^{(t)} + a^{(t)}, \beta_0^{(t)} + b^{(t)})$$

where $a^{(t)}$ is the total number of observations on occasion $t$ ($a^{(t)} = \sum_k \xi_{k,t}$) and $b^{(t)}$ is the number of individuals that were available at occasion $t$.

(4) Sample the identification rate with Gibbs sampling. Similarly to the capture rate, it has a full conditional beta posterior distribution.

$$\alpha|\mathbf{x} \sim \beta(a_0^\alpha + a^\alpha, b_0^\alpha + b^\alpha)$$

where $a^\alpha = \sum_j x_j I(\nu_{j,t} = 1)$ is the total number of correct identifications and $b^\alpha = \sum_j x_j I(\nu_{j,t} = 2)$ the total number of misidentifications.

(5) Sample jointly $N$ and **x**. To update the frequencies of the latent histories **x** and **z** and the number of individuals we use Metropolis-Hastings. Since an individual can be correctly identified on one of its observations and misidentified on another on the same occasion, to propose an $\mathbf{x}'$ we need to be able to add an error to any individual. Except for this point, the concept of the algorithm remains the same. Randomly add or remove a misidentification. To add a misidentification, sample an occasion $t$ from those that could have generated one (*i.e.*, the occasions for which there is at least one individual with a unique capture, at that occasion). Then sample a history $\nu_0$ from all the available ones in the current set of histories. And "merge" a history with a unique capture at occasion $t$ into the sampled history (*i.e.*, add a capture and a misidentification at occasion $t$ to an individual with history $\nu_0$ and remove one history $\nu_{1t}$). To remove a misidentification, sample an occasion $t$ from those where at least one misidentification has occurred. Then sample a history $\nu_2$ that contains a misidentification at that occasion $t$, remove a capture and an error at occasion $t$ to one individual with history $\nu_2$, and add an individual with a unique capture at occasion $t$.

More formally, follow the steps:

(a) Define:
- $\nu_{1t}$ the history with a unique capture at time $t$ (potential ghost),
- $\chi_{2,t}(\mathbf{x}) = \{\nu_j | \nu_{j,t} - o_{j,t} > 0, x_{\nu_j} > 0\}$ the set of histories *containing* at least one ghost at occasion $t$, for the given **x**.

(b) With probability 0.5, go to (i), otherwise go to (ii).
  (i) Add a misidentification (i.e. a ghost) to the latent set.
  - Sample $t \in \{t | x_{\nu_{1t}} > 0\}$.
  - Define $\nu_{1t}$ the latent history with a unique capture at $t$
  - Sample $\nu_0$ uniformly from the set of histories for which $\mathbf{x}_j > 0$.

- Set $\nu_2 = \nu_0 + \nu_{1t}$ and add one error at occasion $t$. For example, if $\nu_{1t} = (0, 1^{(0)})$, and $\nu_0 = (1^{(0)}, 3^{(1)})$, then $\nu_2 = (1^{(0)}, 4^{(2)})$.
- Define the move $b_{\nu_0,\nu_1,\nu_2} = (-1, -1, +1)$.

(ii) Remove a misidentification from the latent set.

- Sample $t$ in the set of occasions where at least one misidentification occurred.
- Sample $\nu_2 \in \chi_{2,t}(x)$, the history containing a misidentification at occasion $t$.
- Define $\nu_0 = \nu_2 - \nu_{1t}$ and remove one error at occasion $t$. For example, if $\nu_{1t} = (0, 1^{(0)})$, and $\nu_2 = (1^{(0)}, 4^{(2)})$ then $\nu_0 = (1^{(0)}, 3^{(1)})$.
- Define the move $b_{\nu_0,\nu_1,\nu_2} = (+1, +1, -1)$.

(c) Define $\mathbf{x}' = \mathbf{x}^{(k-1)} + b$.
(d) Calculate $\mathbf{z}' = \mathbf{B}\mathbf{x}'$ and $N' = \sum \mathbf{x}'$.
(e) With probability $r_1$, set $\mathbf{x}^k = \mathbf{x}'$, $\mathbf{z}^k = \mathbf{z}'$ and $N^(k) = N'$. Otherwise set $\mathbf{x}^k = \mathbf{x}^{k-1}$, $\mathbf{z}^k = \mathbf{z}^{k-1}$ and $N^(k) = N^{(k-1)}$.

(5) $$r_1 = min\left(1, \frac{[\mathbf{y}|\mathbf{x}', \mathbf{z}', N', \boldsymbol{\lambda}, \alpha]}{[\mathbf{y}|\mathbf{x}^{(k-1)}, \mathbf{z}^{(k-1)}, N, \boldsymbol{\lambda}, \alpha]} \frac{q(\mathbf{x}^{(k-1)}|\mathbf{x}')}{q(\mathbf{x}'|\mathbf{x}^{(k-1)})}\right)$$

The proposal densities are calculated by multiplying the probabilities of each sampling step used for defining the move. They are successively: the probability of adding (or removing) an error, the probability of choosing the $\nu_0$ (or $\nu_2$) and the probability of choosing the $t$ knowing the sampled $\nu$. When adding an error, the proposal density $q$ is:

(6) $$q(x'|x^{k-1}) = \frac{0.5}{\#\{t|x_{\nu_{1t}} > 0\}\#\chi_0}$$

and when removing an error, is:

(7) $$q(x'|x^{k-1}) = \frac{0.5}{\#\{t|e_t > 0\}\#\chi_{2t}}$$

where $e_t$ is the number of misidentification at $t$, $\chi_0$ the set of histories with $x_j > 0$ and $\#S$ denotes the cardinality of $S$.

(6) Sample the number of unseen individuals $x_0$:
(a) set $\mathbf{x}' = \mathbf{x}$, $\mathbf{z}' = \mathbf{z}$ and $x_0$ the number of unseen individual in $\mathbf{z}$ (and $\mathbf{x}$),
(b) sample a move $c \in [-D, D]$ where D is fixed integer,
(c) define $x_0' = x_0 + c$,
(d) set the number of unseen individuals in $\mathbf{x}'$ and $\mathbf{z}'$ to $x_0'$,
(e) accept it with probability $r_2$:

(8) $$r_2 = min\left(1, \frac{[\mathbf{z}'|N', \boldsymbol{\lambda}]}{[\mathbf{z}|N, \boldsymbol{\lambda}]}\right).$$

(7) repeat steps 3 to 6 as much as needed.

#### Supplement C: Coverage rate of model $M_{\lambda,\alpha}$ in all tested situations

| S | lambda | alpha | $M_{\lambda,\alpha}$ | $M_t$ | Yoshizaki |
|---|---|---|---|---|---|
| 5 | 0.11 | 0.80 | 7 | 0 | 6 |
| 5 | 0.11 | 0.90 | 8 | 3 | 7 |
| 5 | 0.11 | 0.95 | 7 | 7 | 7 |
| 5 | 0.23 | 0.80 | 9 | 0 | 10 |
| 5 | 0.23 | 0.90 | 10 | 1 | 10 |
| 5 | 0.23 | 0.95 | 8 | 5 | 10 |
| 5 | 0.36 | 0.80 | 8 | 0 | 8 |
| 5 | 0.36 | 0.90 | 9 | 0 | 9 |
| 5 | 0.36 | 0.95 | 9 | 0 | 9 |
| 5 | 0.51 | 0.80 | 10 | 0 | 10 |
| 5 | 0.51 | 0.90 | 10 | 0 | 10 |
| 5 | 0.51 | 0.95 | 10 | 0 | 8 |
| 7 | 0.11 | 0.80 | 2 | 0 | 4 |
| 7 | 0.11 | 0.90 | 6 | 2 | 4 |
| 7 | 0.11 | 0.95 | 6 | 7 | 6 |
| 7 | 0.23 | 0.80 | 9 | 0 | 9 |
| 7 | 0.23 | 0.90 | 10 | 0 | 10 |
| 7 | 0.23 | 0.95 | 10 | 0 | 10 |
| 7 | 0.36 | 0.80 | 9 | 0 | 8 |
| 7 | 0.36 | 0.90 | 10 | 0 | 10 |
| 7 | 0.36 | 0.95 | 9 | 0 | 10 |
| 7 | 0.51 | 0.80 | 10 | 0 | 10 |
| 7 | 0.51 | 0.90 | 10 | 0 | 10 |
| 7 | 0.51 | 0.95 | 10 | 0 | 10 |
| 9 | 0.11 | 0.80 | 7 | 0 | 8 |
| 9 | 0.11 | 0.90 | 9 | 1 | 9 |
| 9 | 0.11 | 0.95 | 9 | 5 | 9 |
| 9 | 0.23 | 0.80 | 10 | 0 | 9 |
| 9 | 0.23 | 0.90 | 10 | 0 | 9 |
| 9 | 0.23 | 0.95 | 10 | 0 | 8 |
| 9 | 0.36 | 0.80 | 10 | 0 | 7 |
| 9 | 0.36 | 0.90 | 10 | 0 | 6 |
| 9 | 0.36 | 0.95 | 10 | 0 | 8 |
| 9 | 0.51 | 0.80 | 9 | 0 | 8 |
| 9 | 0.51 | 0.90 | 10 | 0 | 6 |
| 9 | 0.51 | 0.95 | 10 | 0 | 6 |

#### Supplement D: Capture rate estimates for the otter dataset

| year | Occasion | $M_{\lambda,\alpha}$ | Yoshizaki | $M_t$ |
|---|---|---|---|---|
| | 1 | 1.11 [0.67, 1.67] | 0.66 [0.42, 0.87] | 0.48 [0.29, 0.68] |
| | 2 | 1.52 [0.99, 2.16] | 0.6 [0.35, 0.81] | 0.44 [0.26, 0.64] |
| 2006 | 3 | 1.63 [1.08, 2.29] | 0.72 [0.48, 0.91] | 0.56 [0.37, 0.76] |
| | 4 | 1.64 [1.08, 2.3] | 0.54 [0.29, 0.77] | 0.44 [0.26, 0.64] |
| | 5 | 1.46 [0.94, 2.09] | 0.6 [0.35, 0.82] | 0.48 [0.29, 0.68] |
| | 1 | 0.51 [0.26, 0.84] | 0.46 [0.25, 0.68] | 0.33 [0.18, 0.5] |
| | 2 | 1.48 [1.03, 2.03] | 0.55 [0.34, 0.76] | 0.44 [0.28, 0.62] |
| 2007 | 3 | 1.35 [0.92, 1.87] | 0.7 [0.48, 0.89] | 0.56 [0.39, 0.73] |
| | 4 | 1.44 [0.99, 1.97] | 0.56 [0.34, 0.76] | 0.44 [0.28, 0.62] |
| | 5 | 1.1 [0.72, 1.57] | 0.55 [0.34, 0.77] | 0.39 [0.23, 0.56] |
| | 1 | 1.05 [0.6, 1.63] | 0.73 [0.49, 0.91] | 0.57 [0.37, 0.75] |
| | 2 | 1.11 [0.64, 1.72] | 0.73 [0.49, 0.91] | 0.49 [0.29, 0.68] |
| 2008 | 3 | 1.57 [1, 2.26] | 0.67 [0.43, 0.87] | 0.45 [0.26, 0.64] |
| | 4 | 1.58 [1, 2.27] | 0.67 [0.42, 0.87] | 0.57 [0.37, 0.75] |
| | 5 | 1.31 [0.8, 1.94] | 0.48 [0.26, 0.72] | 0.45 [0.26, 0.64] |
| | 1 | 1.7 [1.14, 2.38] | 0.82 [0.61, 0.96] | 0.73 [0.54, 0.89] |
| | 2 | 1.29 [0.81, 1.88] | 0.64 [0.4, 0.84] | 0.52 [0.32, 0.71] |
| 2010 | 3 | 2.28 [1.62, 3.05] | 0.88 [0.68, 0.99] | 0.73 [0.54, 0.89] |
| | 4 | 1.05 [0.62, 1.6] | 0.76 [0.54, 0.93] | 0.56 [0.36, 0.75] |
| | 5 | 1.58 [1.04, 2.23] | 0.64 [0.4, 0.84] | 0.47 [0.28, 0.67] |
| | 1 | 0.94 [0.59, 1.37] | 0.55 [0.36, 0.74] | 0.49 [0.31, 0.68] |
| | 2 | 0.86 [0.53, 1.27] | 0.6 [0.39, 0.79] | 0.53 [0.34, 0.71] |
| 2011 | 3 | 1.18 [0.78, 1.65] | 0.52 [0.32, 0.71] | 0.53 [0.35, 0.71] |
| | 4 | 1.51 [1.05, 2.04] | 0.63 [0.43, 0.81] | 0.56 [0.38, 0.74] |
| | 5 | 1.35 [0.93, 1.85] | 0.81 [0.62, 0.94] | 0.78 [0.6, 0.91] |
| | 1 | 0.89 [0.52, 1.37] | 0.49 [0.28, 0.7] | 0.38 [0.21, 0.57] |
| | 2 | 1.63 [1.11, 2.25] | 0.74 [0.53, 0.9] | 0.69 [0.5, 0.85] |
| 2012 | 3 | 1.83 [1.28, 2.49] | 0.59 [0.38, 0.79] | 0.5 [0.31, 0.68] |
| | 4 | 2.36 [1.72, 3.09] | 0.94 [0.8, 1] | 0.76 [0.58, 0.9] |
| | 5 | 1.89 [1.32, 2.56] | 0.74 [0.52, 0.9] | 0.61 [0.42, 0.78] |

#### Supplement E: Simulation study with small population size

In order to known how the $M_{\lambda,\alpha}$ model performs in situations similar to the otter dataset, we simulated 50 datasets with a population size $N = 30$, a capture rate $\lambda = 0.5$ and identification rate $\alpha = 0.95$. This is close to what the $M_{\lambda,\alpha}$ model estimates for the otter dataset. We estimated the population size using the $M_{\lambda,\alpha}$ model, with uninformative priors as it was done in the main paper study.

The population size estimates are shown in Figure 1. There is an average bias of $-1.8$ ($\approx -6\%$) in the estimates of the population size. However this bias is not systematic and the true population size is most of the time in the 95% credible interval.

**Figure 1** – Small population simulation study, population size estimates. The horizontal line is the true population size. The red points are those for which the true population size is not in the 95%CI (error bars)

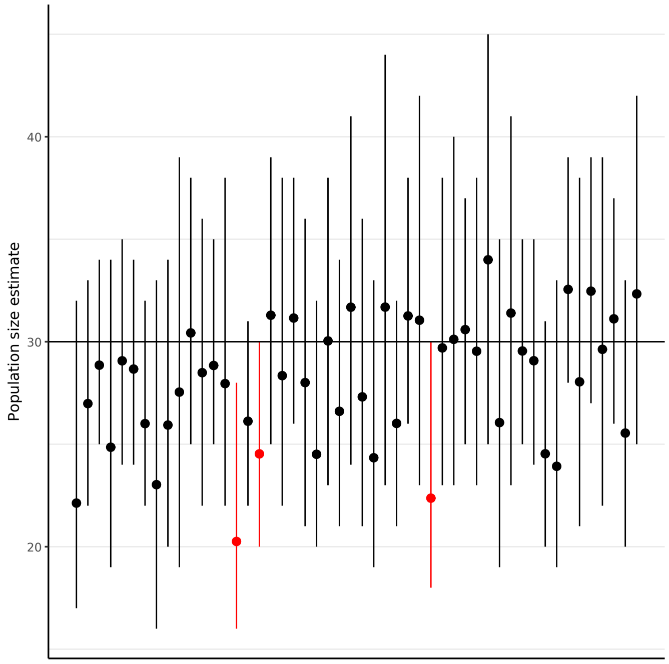


**Figure 2** – Small population simulation study, $\lambda_t$ estimates. There are five estimates per simulation (close to each other) for the five capture occasions. The horizontal line is the true simulated $\lambda$. The red points are those for which the true population size is not in the 95%CI (error bars), the blue points are those for which the true $\lambda$ is not in the 95%CI, the purple points are those for which neither the true population size and the true $\lambda$ are in the 95%CI.

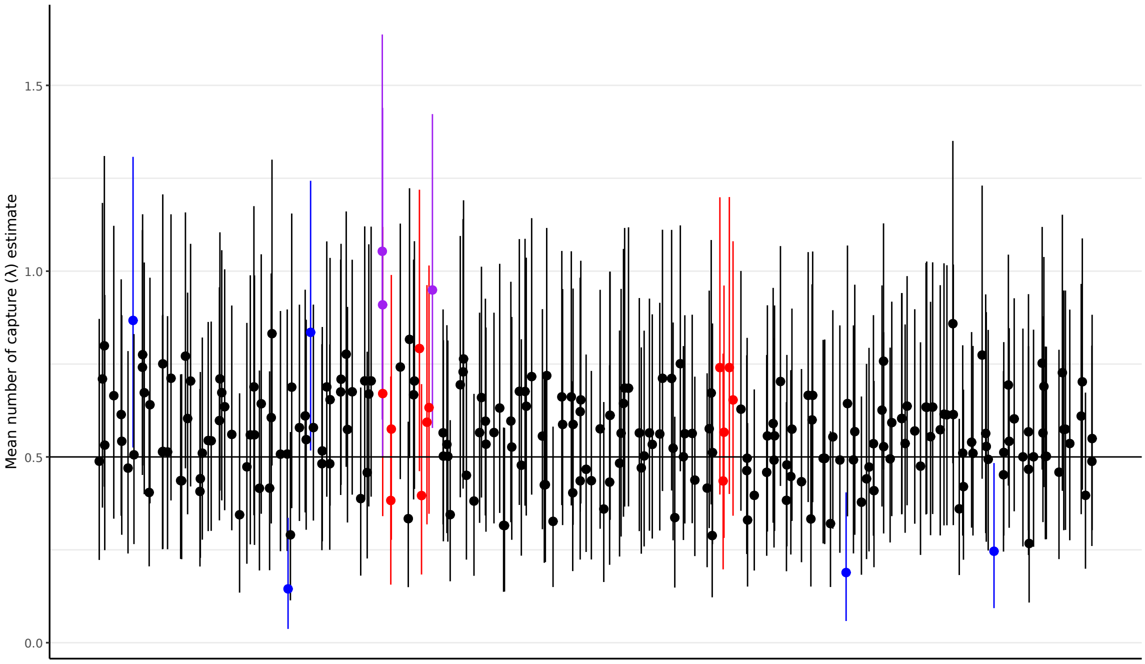


The estimated lambda are shown in Figure 2. There is a small average bias but most true values are in the 95%CI. In addition, for two of the three simulations for which the true population size was not in the 95%CI, one or two $\lambda_t$ are highly overestimated (estimated two times the real value) but for the third simulation with underestimated *N*, the $\lambda_t$ are correctly estimated.

The estimated $\alpha$ are shown on Figure 3. Again, there is a small average bias but not systematic and most true values fall in the 95%CI.

**Figure 3** – Small population simulation study, population size estimates. The horizontal line is the true population size. The red points are those for which the true population size is not in the 95%CI (error bars)

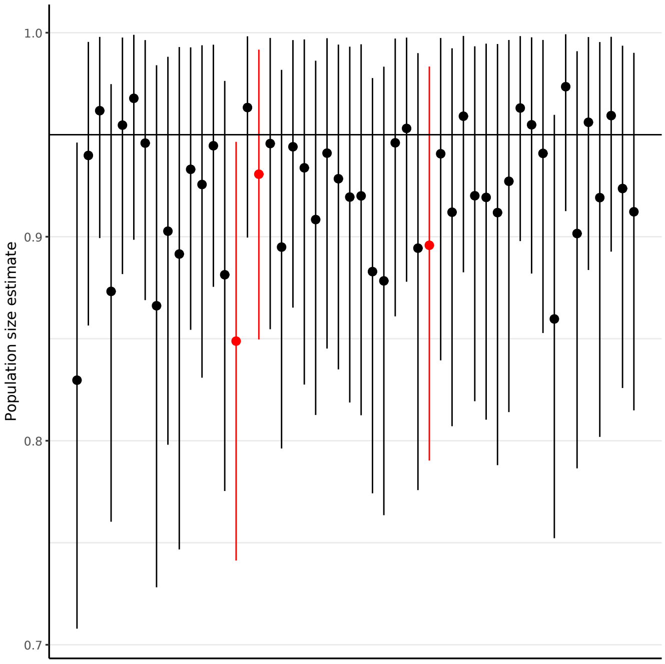

team 4 (n.d.). *4ti2—A software package for algebraic, geometric and combinatorial problems on linear spaces*. URL: https://4ti2.github.io.
